# An analytic approach for interpretable predictive models in high dimensional data, in the presence of interactions with exposures

**DOI:** 10.1101/102475

**Authors:** Sahir Rai Bhatnagar, Yi Yang, Budhachandra Khundrakpam, Alan C Evans, Mathieu Blanchette, Luigi Bouchard, Celia MT Greenwood

**Affiliations:** Department of Epidemiology, Biostatistics and Occupational Health, McGill University Lady Davis Institute, Jewish General Hospital, Montréal, QC; Department of Mathematics and Statistics, McGill University; Montreal Neurological Institute, McGill University; Department of Computer Science, McGill University; Department of Biochemistry, Université de Sherbrooke

**Keywords:** gene-environment interaction, high-dimensional clustering, prediction models, topological overlap matrix, penalized regression

## Abstract

Predicting a phenotype and understanding which variables improve that prediction are two very challenging and overlapping problems in analysis of high-dimensional data such as those arising from genomic and brain imaging studies. It is often believed that the number of truly important predictors is small relative to the total number of variables, making computational approaches to variable selection and dimension reduction extremely important. To reduce dimensionality, commonly-used two-step methods first cluster the data in some way, and build models using cluster summaries to predict the phenotype.

It is known that important exposure variables can alter correlation patterns between clusters of high-dimensional variables, i.e., alter network properties of the variables. However, it is not well understood whether such altered clustering is informative in prediction. Here, assuming there is a binary exposure with such network-altering effects, we explore whether use of exposure-dependent clustering relationships in dimension reduction can improve predictive modelling in a two-step framework. Hence, we propose a modelling framework called ECLUST to test this hypothesis, and evaluate its performance through extensive simulations.

With ECLUST, we found improved prediction and variable selection performance compared to methods that do not consider the environment in the clustering step, or to methods that use the original data as features. We further illustrate this modelling framework through the analysis of three data sets from very different fields, each with high dimensional data, a binary exposure, and a phenotype of interest. Our method is available in the *eclust* CRAN package.

## Introduction

In this article, we consider the prediction of an outcome variable *y* observed on *n* individuals from *p* variables, where *p* is much larger than *n*. Challenges in this high-dimensional context include not only building a good predictor which will perform well in an independent dataset, but also being able to interpret the factors that contribute to the predictions. This latter issue can be very challenging in ultra-high dimensional predictor sets. For example, multiple different sets of covariates may provide equivalent measures of goodness of fit (Fan et al., 2014), and therefore how does one decide which are important? If many variables are highly correlated, interpretation may be improved by acknowledging the existence of an underlying or latent factor generating these patterns. In consequence, many authors have suggested a two-step procedure where the first step is to cluster or group variables in the design matrix in an interpretable way, and then to perform model fitting in the second step using a summary measure of each group of variables.

There are several advantages to these two-step methods. Through the reduction of the dimension of the model, the results are often more stable with smaller prediction variance, and through identification of sets of correlated variables, the resulting clusters can provide an easier route to interpretation. From a practical point of view, two-step approaches are both flexible and easy to implement because efficient algorithms exist for both clustering (e.g. (Müllner, 2013)) and model fitting (e.g. (Friedman et al., 2010; Yang and Zou, 2014; Kuhn, 2008)), particularly in the case when the outcome variable is continuous.

This two-step idea dates back to 1957 when Kendall first proposed using principal components in regression (Kendall, 1957). Hierarchical clustering based on the correlation of the design matrix has also been used to create groups of genes in microarray studies. For example, at each level of a hierarchy, cluster averages have been used as new sets of potential predictors in both forward-backward selection (Hastie et al., 2001) or the lasso (Park et al., 2007). Bühlmann *et al.* proposed a bottom-up agglomerative clustering algorithm based on canonical correlations and used the group lasso on the derived clusters (Bühlmann et al., 2013). A more recent proposal performs sparse regression on cluster prototypes (Reid and Tibshirani, 2016), i.e., extracting the most representative gene in a cluster instead of averaging them.

These two-step approaches usually group variables based on a matrix of correlations or some transformation of the correlations. However, when there are external factors, such as exposures, that can alter correlation patterns, a dimension reduction step that ignores this information may be suboptimal. Many of the high-dimensional genomic data sets currently being generated capture a possibly dynamic view of how a tissue is functioning, and demonstrate differential patterns of coregulation or correlation under different conditions. We illustrate this critical point with an example of a microarray gene expression dataset available in the *COPDSexualDimorphism.data* package (Sathirapongsasuti, 2013) from Bioconductor. This study measured gene expression in Chronic Obstructive Pulmonary Disease (COPD) patients and controls in addition to their age, gender and smoking status. To see if there was any effect of smoking status on gene expression, we plotted the expression profiles separately for current and never smokers. To balance the covariate profiles, we matched subjects from each group on age, gender and COPD case status, resulting in a sample size of 7 in each group. Heatmaps in Figure 1 show gene expression levels and the corresponding gene-gene correlation matrices as a function of dichotomized smoking status for 2,900 genes with large variability. Evidently, there are substantial differences in correlation patterns between the smoking groups (Figures 1a and 1b). However, it is difficult to discern any patterns or major differences between the groups when examining the gene expression levels directly (Figures 1c and 1d). This example highlights two key points; 1) environmental exposures can have a widespread effect on regulatory networks and 2) this effect may be more easily discerned by looking at a measure for gene similarity, relative to analyzing raw expression data. Many other examples of altered co-regulation and phenotype associations can be found. For instance, in a pediatric brain development study, very different correlation patterns of cortical thickness within brain regions were observed across age groups, consistent with a process of fine-tuning an immature brain system into a mature one (Khundrakpam et al., 2013). A comparison of gene expression levels in bone marrow from 327 children with acute leukemia found several differentially *coexpressed* genes in philadelphia positive leukemias compared to the cytogenetically normal group (Kostka and Spang, 2004). To give a third example, an analysis of RNA-sequencing data from The Cancer Genome Atlas (TCGA) revealed very different correlation patterns among sets of genes in tumors grouped according to their missense or null mutations in the TP53 tumor suppressor gene (Oros Klein et al., 2016).

**Figure 1.**
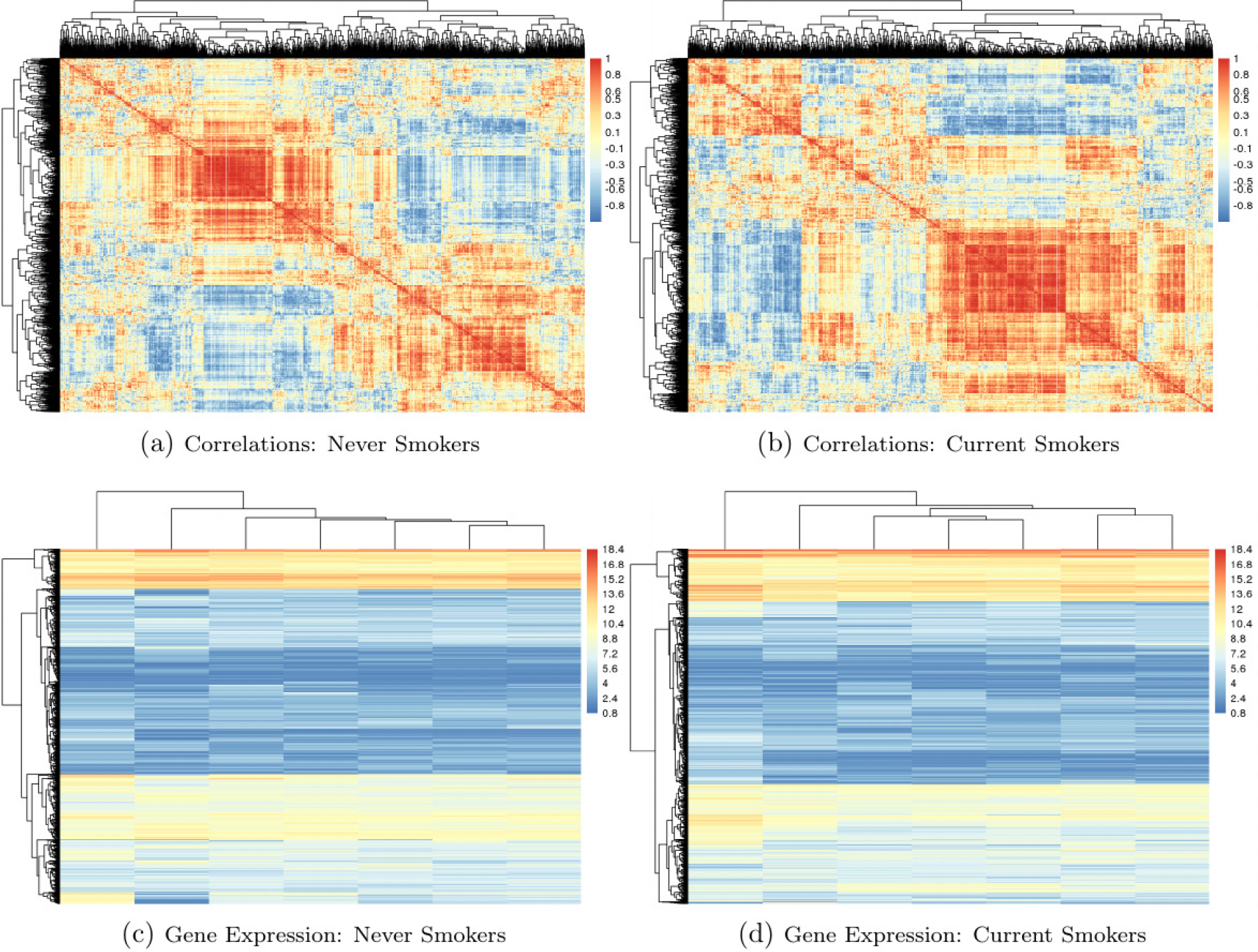
Heatmaps of correlations between genes (top) and gene expression data (bottom - rows are genes and columns are subjects), stratified by smoking status from a microarray study of COPD (Sathirapongsasuti, 2013). The 20% most variable genes are displayed (2,900 genes). There are 7 subjects in each group, matched on COPD case status, gender and age. Data available on Bioconductor in the `COPDSexualDimorphism.data` package.

Therefore, in this paper, we pose the question whether clustering or dimension reduction that incorporates known covariate or exposure information can improve prediction models in high dimensional genomic data settings. Substantial evidence of dysregulation of genomic coregulation has been observed in a variety of contexts, however we are not aware of any work that carefully examines how this might impact the performance of prediction models. We propose a conceptual analytic strategy called ECLUST, for prediction of a continuous or binary outcome in high dimensional contexts while exploiting exposure-sensitive data clusters. We restrict our attention to two-step algorithms in order to implement a covariate-driven clustering.

Specifically, we hypothesize that within two-step methods, variable grouping that considers exposure information can lead to improved predictive accuracy and interpretability. We use simulations to compare our proposed method to comparable approaches that combine data reduction with predictive modelling. We are focusing our attention primarily on the performance of alternative dimension reduction strategies within the first step of a two-step method. Therefore, performance of each strategy is compared for several appropriate step 2 predictive models. We then illustrate these concepts more concretely by analyzing three data sets. Our method and the functions used to conduct the simulation studies have been implemented in the **R** package **eclust** (Bhatnagar, 2017), available on CRAN. Extensive documentation of the package is available at http://sahirbhatnagar.com/eclust/.

## Methods

Assume there is a single binary environmental factor *E* of importance, and an *n* × *p* high dimensional (HD) data set **X** (*n* observations, *p* features) of relevance. This could be genome-wide epigenetic data, gene expression data, or brain imaging data, for example. Assume there is a continuous or binary phenotype of interest *Y* and that the environment has a widespread effect on the HD data, i.e., affects many elements of the HD data. The primary goal is to improve prediction of *Y* by identifying interactions between *E* and **X** through a carefully constructed data reduction strategy that exploits *E* dependent correlation patterns. The secondary goal is to improve identification of the elements of **X** that are involved; we denote this subset by *S*_0_. We hypothesize that a systems-based perspective will be informative when exploring the factors that are associated with a phenotype of interest, and in particular we hypothesize that incorporation of environmental factors into predictive models in a way that retains a high dimensional perspective will improve results and interpretation.

### Potential impacts of covariate-dependent coregulation

Motivated by real world examples of differential coexpression, we first demonstrate that environment-dependent correlations in **X** can induce an interaction model. Without loss of generality, let *p* = 2 and the relationship between *X*_1_ and *X*_2_ depend on the environment such that

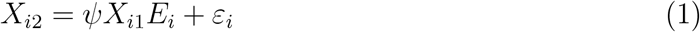

where *ɛ*_*i*_ is an error term and *ψ* is a slope parameter, that is:

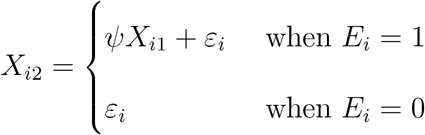

Consider the 3-predictor regression model

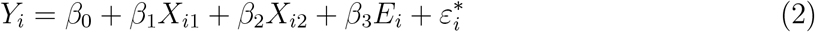

where 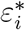 is another error term which is independent of *ɛ*_*i*_. At first glance (2) does not contain any interaction terms. However, substituting (1) for *X*_*i*2_ in (2) we get

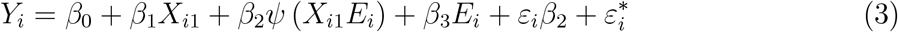

The third term in (3) resembles an interaction model, with *β_2_ψ* being the interaction parameter. We present a second illustration showing how non-linearity can induce interactions. Suppose

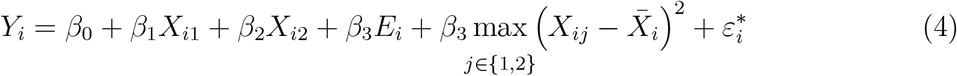

Substituting (1) for *X*_*i*2_ in (4) we obtain a non-linear interaction term. Equation (4) **provided partial motivation for the model used in our third simulation scenario.** Some motivation for this model and a graphical representation are presented below in the Simulation Studies section.

### Proposed framework and algorithm

We restrict attention to methods containing two phases as illustrated in Figure 2: 1a) a clustering stage where variables are clustered based on some measure of similarity, 1b) a dimension reduction stage where a summary measure is created for each of the clusters, and 2) a simultaneous variable selection and regression stage on the summarized cluster measures. Although this framework appears very similar to any two-step approach, our hypothesis is that allowing the clustering in Step 1a to depend on the environment variable can lead to improvements in prediction after Step 2. Hence, methods in Step 1a are adapted to this end, as decribed in the following sections. Our focus in this manuscript is on the clustering and cluster representation steps. Therefore, we compare several well known methods for variable selection and regression that are best adapted to our simulation designs and data sets.

**Figure 2.**
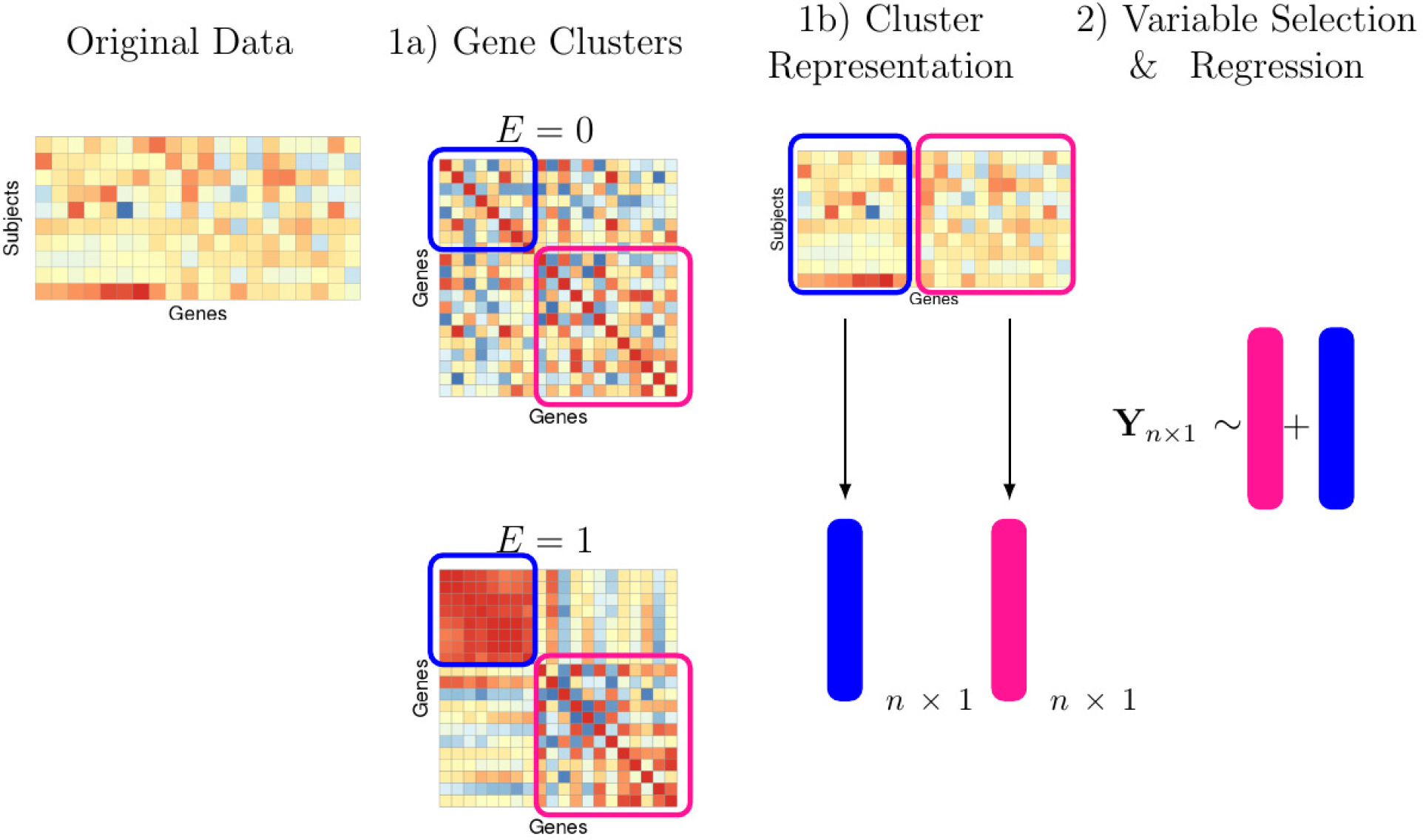
Overview of our proposed method. 1a) A measure of similarity is calculated separately for both groups and clustering is performed on a linear combination of these two matrices. 1b) We reduce the dimension of each cluster by taking a summary measure. 2) Variable selection and regression is performed on the cluster representatives, E and their interaction with E.

**Step 1a: Clustering using co-expression networks that are influenced by the environment.** In agglomerative clustering, a measure of similarity between sets of observations is required in order to decide which clusters should be combined. Common choices include Euclidean, maximum and absolute distance. A more natural choice in genomic or brain imaging data is to use Pearson correlation (or its absolute value) because the derived clusters are biologically interpretable. Indeed, genes that cluster together are correlated and thus likely to be involved in the same cellular process. Similarly, cortical thickness measures of the brain tend to be correlated within pre-defined regions such as the left and right hemisphere, or frontal and temporal regions (Sato et al., 2013). However, the information on the connection between two variables, as measured by the Pearson correlation for example, may be noisy or incomplete. Thus it is of interest to consider alternative measures of pairwise interconnectedness. Gene co-expression networks are being used to explore the system-level function of genes, where nodes represent genes and are connected if they are significantly co-expressed (Zhang and Horvath, 2005), and here we use their overlap measure (Ravasz et al., 2002) to capture connectnedness between two *X* variables within each environmental condition. As was discussed earlier, genes can exhibit very different patterns of correlation in one environment versus the other (e.g. Figure 1). Furthermore, measures of similarity that go beyond pairwise correlations and consider the shared connectedness between nodes can be useful in elucidating networks that are biologically meaningful. Therefore, we propose to first look at the topological overlap matrix (TOM) separately for exposed (*E* = 1) and unexposed (*E* = 0) individuals (**see Supplemental Section A for details on the TOM**). We then seek to identify nodes that are very different between environments. We determine differential coexpression using the absolute difference *TOM*(*X*_diff_) = |*TOM*_*E*=1_ – *TOM*_*E*=0_| (Oros Klein et al., 2016). We then use hierarchical clustering with average linkage on the derived difference matrix to identify these differentially co-expressed variables. Clusters are automatically chosen using the `dynamicTreeCut` (Langfelder et al., 2008) algorithm. Of course, there could be other clusters which are not sensitive to the environment. For this reason we also create a set of clusters based on the TOM for all subjects denoted *TOM*(*X*_all_). This will lead to each covariate appearing in two clusters. In the sequel we denote the clusters derived from *TOM*(*X*_all_) as the set *C*_all_ = {*C*_1_,…,*C*_*k*_}, and those derived from *TOM*(*X*_diff_) as the set *C*_diff_ = {*C*_*k*+1_,…,*C*_*l*_} where *k* < *ℓ* < *p*.

**Step 1b: Dimension reduction via cluster representative.** Once the clusters have been identified in phase 1, we proceed to reduce the dimensionality of the overall problem by creating a summary measure for each cluster. A low-dimensional structure, i.e. grouping when captured in a regression model, improves predictive performance and facilitates a model’s interpretability. We propose to summarize a cluster by a single representative number. Specifically, we chose the average values across all measures (Park et al., 2007; Bühlmann et al., 2013), and the first principal component (Langfelder and Horvath, 2007). These representative measures are indexed by their cluster, i.e., the variables to be used in our predictive models are 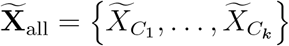 for clusters that do not consider *E*, as well as 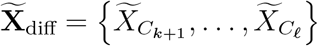 for *E*-derived clusters. The tilde notation on the *X* is to emphasize that these variables are different from the separate variables in the original data.

**Step 2: Variable Selection and Regression.** Because the clustering in phase 1 is unsupervised, it is possible that the derived latent representations from phase 2 will not be associated with the response. We therefore use penalized methods for supervised variable selection, including the lasso (Tibshirani, 1996) and elasticnet (Zou and Hastie, 2005) for linear models, and multivariate adaptive regression splines (MARS) (Friedman,1991) for nonlinear models. We argue that the selected non-zero predictors in this model will represent clusters of genes that interact with the environment and are associated with the phenotype. Such an additive model might be insufficient for predicting the outcome. In this case we may directly include the environment variable, the summary measures and their interaction. In the light of our goals to improve prediction and interpretability, we consider the following model

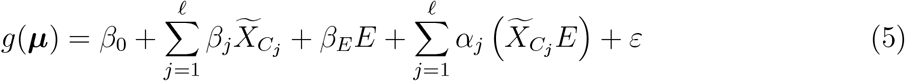

where *g*(·) is a known link function, ***μ*** = **E**[*Y* |**X**, *E*, *β*,*α*] and 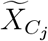 are linear combinations of **X** (from Step 1b). The primary comparison is models with 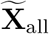 only versus models with 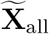 and 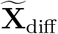. Given the context of either the simulation or the data set, we use either linear models or non linear models. Our general approach, ECLUST, can therefore be summarized by the algorithm in Table 1.

**Table 1.** Details of ECLUST algorithm

| Step | Description, Software<sup>a</sup> and Reference |
| --- | --- |
| 1a) | i) Calculate $TOM$ separately for observations with $E = 0$ and $E = 1$ using `WGCNA::TOMsimilarityFromExpr` (Langfelder and Horvath, 2008)<br>ii) Euclidean distance matrix of $\vert TOM_{E=1} - TOM_{E=0}\vert $ using `stats::dist`<br>iii) Run the `dynamicTreeCut` algorithm (Langfelder et al., 2008, 2016) on the distance matrix to determine the number of clusters and cluster membership using `dynamicTreeCut::cutreeDynamic` with `minClusterSize = 50` |
| 1b) | i) 1st PC or average for each cluster using `stat::prcomp` or `base::mean`<br>ii) Penalized regression model: create a design matrix of the derived cluster representatives and their interactions with $E$ using `stats::model.matrix`<br>iii) MARS model: create a design matrix of the derived cluster representatives and $E$ |
| 2) | i) For linear models, run penalized regression on design matrix from step 1b) using `glmnet::cv.glmnet` (Friedman et al., 2010). Elasticnet mixing parameter `alpha=1` corresponds to the lasso and `alpha=0.5` corresponds to the value we used in our simulations for elasticnet. The tuning parameter `alpha` is selected by minimizing 10 fold cross-validated mean squared error (MSE).<br>ii) For non-linear effects, run MARS on the design matrix from step 1b) using `earth::earth` (Milborrow. Derived from mda:mars by T. Hastie and R. Tibshirani., 2011) with pruning method `pmethod = "backward"` and maximum number of model terms `nk = 1000`. The `degree=1,2` is chosen using 10 fold cross validation (CV), and within each fold the number of terms in the model is the one that minimizes the generalized cross validated (GCV) error. |

<sup>a</sup>All functions are implemented in R (R Core Team, 2016). The naming convention is as follows:
`package_name::package_function`. Default settings used for all functions unless indicated otherwise.

## Simulation Studies

We have evaluated the performance of our ECLUST method in a variety of simulated scenarios. For each simulation scenario we compared ECLUST to the following analytic approaches 1) regression and variable selection is performed on the model which consists of the original variables, *E* and their interaction with *E* (SEPARATE), and 2) clustering is performed without considering the environmental exposure followed by regression and variable selection on the cluster representations, *E*, and their interaction with *E* (CLUST). A detailed description of the methods being compared is summarized in Table 2. We have designed 6 simulation scenarios that illustrate different kinds of relationships between the variables and the response. For all scenarios, we have created high dimensional data sets with *p* predictors (*p* = 5000), and sample sizes of *n* = 200. We also assume that we have two data sets for each simulation - a training data set where the parameters are estimated, and a testing data set where prediction performance is evaluated, each of size *n*_*train*_ = *n*_*test*_ = 200. The number of subjects who were exposed (*n*_*E*=1_ = 100) and unexposed (*n*_*E*=0_ = 100) and the number of truly associated parameters (|*S*_0_| = 500) remain fixed across the 6 simulation scenarios. Let

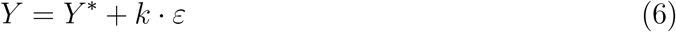

where *Y*\* is the linear predictor, the error term ε is generated from a standard normal distribution, and *k* is chosen such that the signal-to-noise ratio *SNR* = (*Var*(*Y*\*)/*Var*(*ε*)) is 0.2, 1 and 2 (e.g. the variance of the response variable *Y* due to *ε* is 1/*SNR* of the variance of *Y* due to *Y*\*).

**Table 2.**
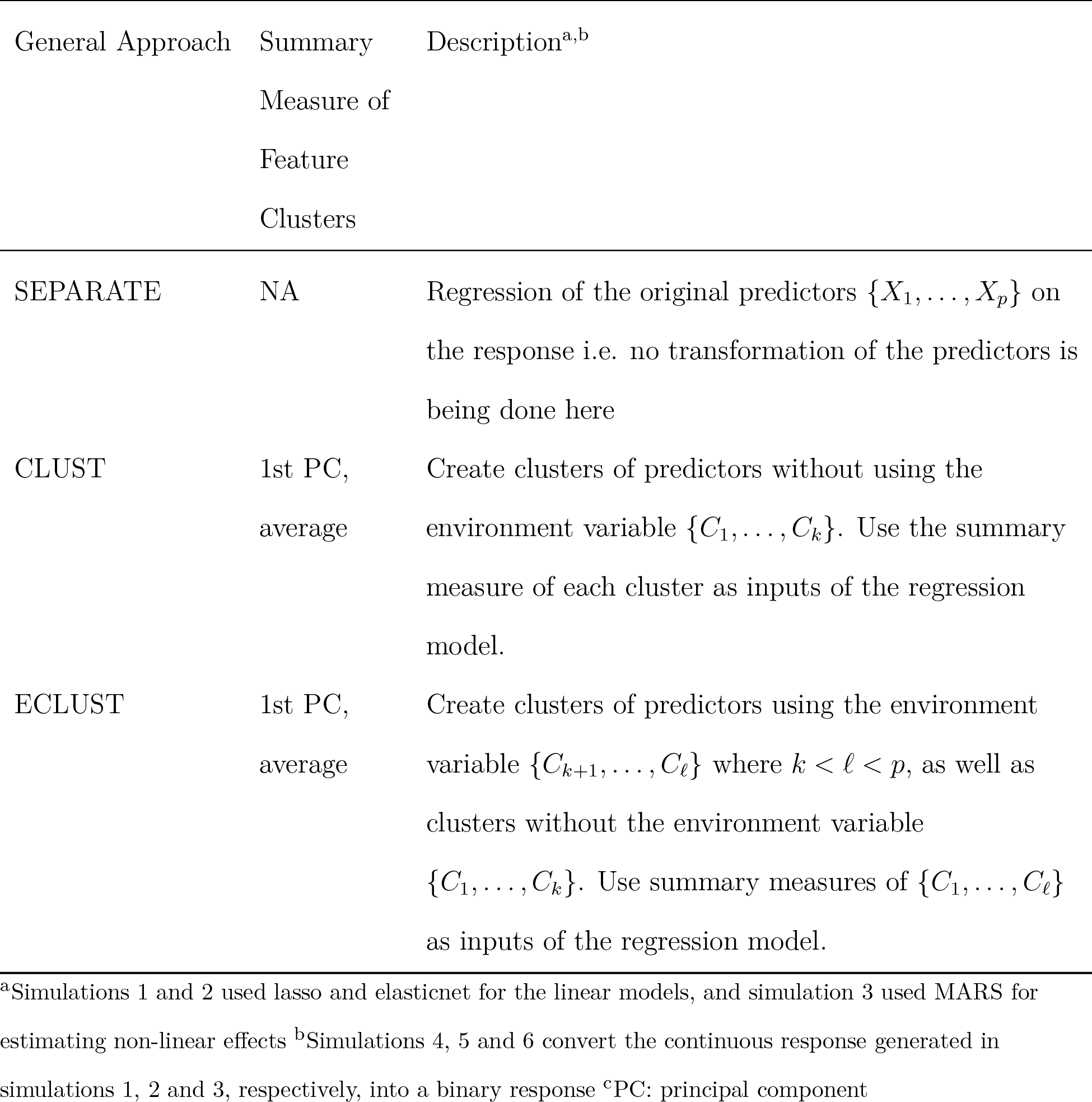
Summary of methods used in simulation study

### The Design Matrix

We generated covariate data in blocks using the `simulateDatExpr` function from the `WGCNA` package in `R` (version 1.51). This generates data from a latent vector: first a seed vector is simulated, then covariates are generated with varying degree of correlation with the seed vector in a given block. We simulated five clusters (blocks), each of size 750 variables, and labeled them by colour (turquoise, blue, red, green and yellow), while the remaining 1250 variables were simulated as independent standard normal vectors (grey) (Figure 3). For the unexposed observations (*E* = 0), only the predictors in the yellow block were simulated with correlation, while all other covariates were independent within and between blocks. The TOM values are very small for the yellow cluster because it is not correlated with any of its neighbors. For the exposed observations (*E* = 1), all 5 blocks contained predictors that are correlated. The blue and turquoise blocks are set to have an average correlation of 0.6. The average correlation was varied for both green and red clusters *ρ* = {0.2, 0.9} and the active set *S*_0_, that are directly associated with *y*, was distributed evenly between these two blocks. Heatmaps of the TOM for this environment dependent correlation structure are shown in Figure 3 with annotations for the true clusters and active variables. This design matrix shows widespread changes in gene networks in the exposed environment, and this subsequently affects the phenotype through the two associated clusters. There are also pathways that respond to changes in the environment but are not associated with the response (blue and turquoise), while others that are neither active in the disease nor affected by the environment (yellow).

**Figure 3.**
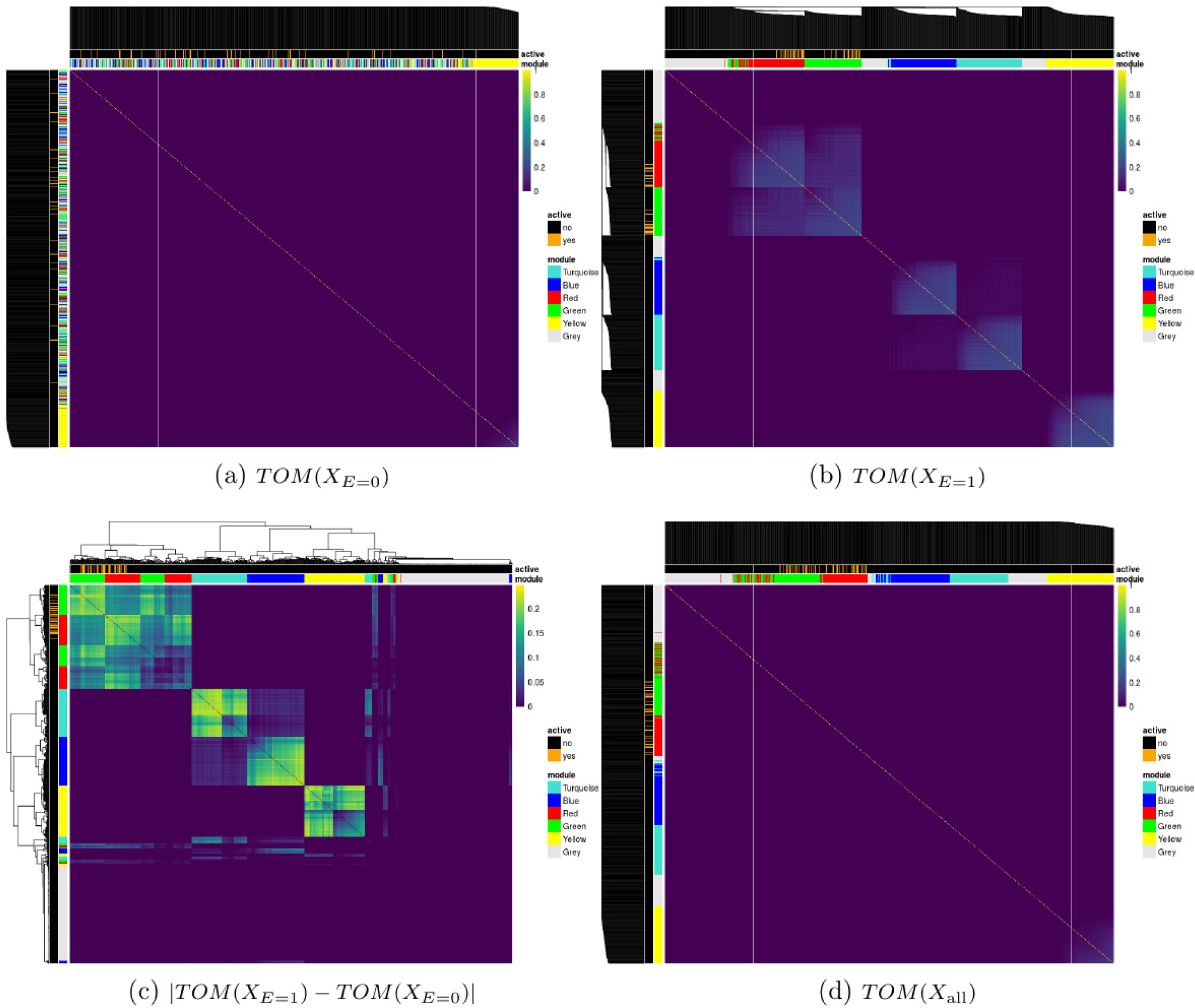
Topological overlap matrices (TOM) of simulated predictors based on subjects with (a) *E* = 0, (b) *E* = 1, (c) their absolute difference and (d) all subjects. Dendrograms are from hierarchical clustering (average linkage) of one minus the TOM for a, b, and d and the euclidean distance for c. Some variables in the red and green clusters are associated with the outcome variable. The *module* annotation represents the true cluster membership for each predictor, and the *active* annotation represents the truly associated predictors with the response.

### The response

The first three simulation scenarios differ in how the linear predictor *Y*\* in (6) is defined, and also in the choice of regression model used to fit the data. In simulations 1 and 2 we use lasso (Tibshirani, 1996) and elasticnet (Zou and Hastie, 2005) to fit linear models; then we use MARS (Friedman, 1991) in simulation 3 to estimate non-linear effects. Simulations 4, 5 and 6 use the GLM version of these models, respectively, since the responses are binary.

### Simulation 1

Simulation 1 was designed to evaluate performance when there are no explicit interactions between X and E (see Equation (3)). We generated the linear predictor from

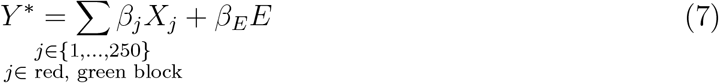

where *β*_*j*_ ~ Unif [0.9,1.1] and *β*_*E*_ = 2. That is, only the first 250 predictors of both the red and green blocks are active. In this setting, only the main effects model is being fit to the simulated data.

### Simulation 2

In the second scenario we explicitly simulated interactions. All non-zero main effects also had a corresponding non-zero interaction effect with *E*. We generated the linear predictor from

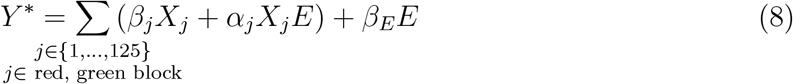

where *β*_*j*_ ~ Unif [0.9,1.1], *α*_*j*_ ~ Unif [0.4,0.6] or *α*_*j*_ ~ Unif [1.9, 2.1], and *β*_*E*_ = 2. In this setting, both the main effects and their interactions with E are being fit to the simulated data.

### Simulation 3

In the third simulation we investigated the performance of the ECLUST approach in the presence of non-linear effects of the predictors on the phenotype:

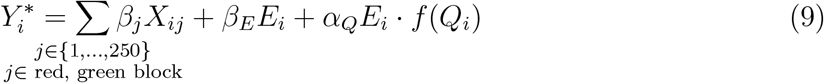

where

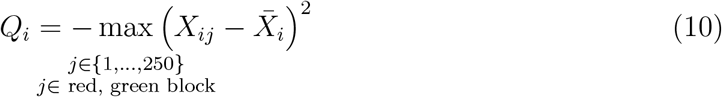

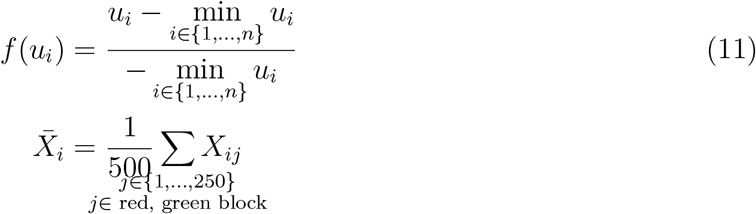

**The design of this simulation was partially motivated by considering the idea of canalization, where systems operate within appropriate parameters until sufficient perturbations accumulate (e.g. Gibson (2009))**. In this third simulation, we set *β*_*j*_ ~ Unif [0.9,1.1], *β*_*E*_ = 2 and *α*_*Q*_ = 1. We assume the data has been appropriately normalized, and that the correlation between any two features is greater than or equal to 0. In simulation 3, we tried to capture the idea that an exposure could lead to coregulation or disregulation of a cluster of *X*’s, which in itself directly impacts Y. Hence, we defined coregulation as the *X*’s being similar in magnitude and disregulation as the *X*’s being very different. The *Q*_*i*_ term in (10) is defined such that higher values would correspond to strong coregulation whereas lower values correspond to disregulation. For example, suppose *Q*_*i*_ ranges from −5 to 0. It will be −5 when there is lots of variability (disregulation) and 0 when there is none (strong coregulation). The function *f* (·) in (11) simply maps *Q*_*i*_ to the [0,1] range. In order to get an idea of the relationship in (9), Figure 4 displays the response *Y* as a function of the first principal component of Σ_*j*_*β*_*j*_*X*_*ij*_ (denoted by `1st PC`) and *f*(*Q*_*i*_). We see that lower values of *f* (*Q*_*i*_) (which implies disregulation of the features) leads to a lower *Y*. In this setting, although the clusters do not explicitly include interactions between the *X* variables, the MARS algorithm allows for the possibility of two way interactions between any of the variables.

**Figure 4.**
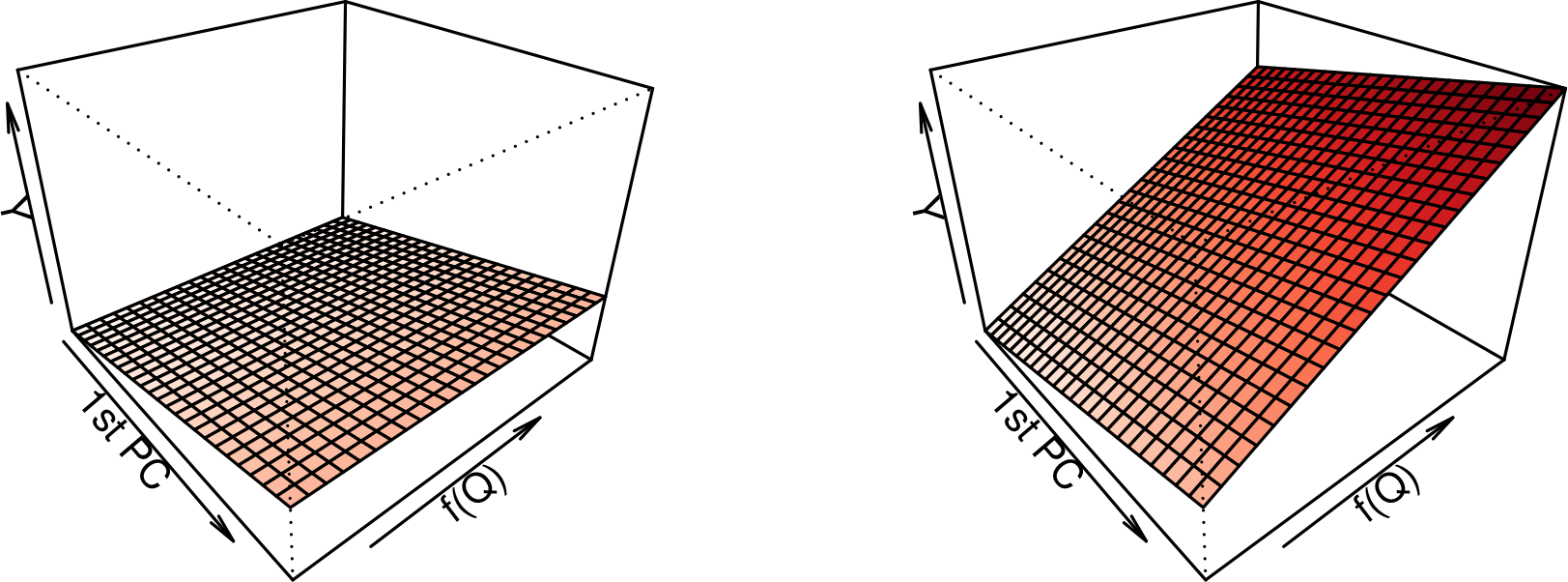
Visualization of the relationship between the response, the first principal component of the main effects and *f* (*Q*_*i*_) in (9) for *E* = 0 (left) and *E* =1 (right) in simulation scenario 3. This graphic also depicts the intuition behind model (4).

### Simulation 4, 5 and 6

We used the same simulation setup as above, except that we took the continuous outcome *Y*, defined *p* = 1/(1 + *exp*(–*Y*)) and used this to generate a two-class outcome *z* with Pr(*z* = 1) = *p* and Pr(*z* = 0) = 1 – *p*. The true parameters were simulated as *β*_*j*_ ~ Unif[log(0.9),log(1.1)], *β*_*E*_ = log(2), *α*_*j*_ ~ Unif[log(0.4),log(0.6)] or *α*_*j*_ ~ Unif[log(1.9),log(2.1)]. Simulations 4, 5 and 6 are the binary response versions of simulations 1, 2 and 3, respectively.

### Measures of Performance

Simulation performance was assessed with measures of model fit, prediction accuracy and feature stability. Several measures for each of these categories, and the specific formulae used are provided in Table 3. We simulated both a training data set and a test data set for each simulation: all tuning parameters for model selection were selected using the training sets only. Although most of the measures of model fit were calculated on the test data sets, true positive rate, false positive rate and correct sparsity were calculated on the training set only. The root mean squared error is determined by predicting the response for the test set using the fitted model on the training set. The area under the curve is determined using the trapezoidal rule (Robin et al., 2011). The stability of feature importance is defined as the variability of feature weights under perturbations of the training set, i.e., small modifications in the training set should not lead to considerable changes in the set of important covariates (Toloşi and Lengauer, 2011). A feature selection algorithm produces a weight (e.g. *β* = (*β*_1_,…,*β*_*p*_)), a ranking (e.g. *rank*(*β*):**r** = (*r*_1_,…,*r*_*m*_)) and a subset of features (e.g. **s** = (*s*_1_,…,*s*_*p*_), *S*_j_ = 𝕀 {*β*_*j*_ ≠ 0} where 𝕀 {·} is the indicator function). In the CLUST and ECLUST methods, we defined a predictor to be non-zero if its corresponding cluster representative weight was non-zero. Using 10-fold cross validation (CV), we evaluated the similarity between two features and their rankings using Pearson and Spearman correlation, respectively. For each CV fold we re-ran the models and took the average Pearson/Spearman correlations of the 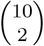 combinations of estimated coefficients vectors. To measure the similarity between two subsets of features we took the average of the Jaccard distance in each fold. A Jaccard distance of 1 indicates perfect agreement between two sets while no agreement will result in a distance of 0. For MARS models we do not report the Pearson/Spearman stability rankings due to the adaptive and functional nature of the model (there are many possible combinations of predictors, each of which are linear basis functions).

**Table 3.** Measures of Performance

| Measure | Formula |
| --- | --- |
| *Model Fit* |  |
| True Positive Rate (TPR) | $\vert \widehat{S} \in S_0\vert /\vert S_0\vert $ |
| False Positive Rate (TPR) | $\vert \widehat{S} \notin S_0\vert /\vert j \notin S_0\vert $ |
| Correct Sparsity (Witten et al., 2014) | $\frac{1}{p} \sum_{j=1}^p A_j$ $A_j = \begin{cases} 1 & \text{if } \hat{\beta}_j = \beta_j = 0 \\ 1 & \text{if } \hat{\beta}_j \neq 0, \beta_j \neq 0 \\ 0 & \text{if else} \end{cases}$ |
| *Prediction Accuracy* |  |
| Root Mean Squared Error (RMSE) | $\ \mathbf{Y}_{test} - \hat{\boldsymbol{\mu}}(\mathbf{X}_{test})\ _2$ |
| Area Under the Curve (AUC) | Trapezoidal rule |
| Hosmer-Lemeshow Test ($G = 10$) | $\chi^2$ test statistic |
| *Feature Stability using K-fold Cross-Validation on training set (Kalousis et al., 2007)* |  |
| Pearson Correlation ($\rho$) (Pearson, 1895) | $\binom{K}{2}^{-1} \sum_{i,j \in \{1, \dots, K\}, i \neq j} \frac{cov(\hat{\beta}_{(i)}, \hat{\beta}_{(j)})}{\sigma_{\hat{\beta}_{(i)}} \sigma_{\hat{\beta}_{(j)}}}$ |
| Spearman Correlation ($r$) (Spearman, 1904) | $\binom{K}{2}^{-1} \sum_{i,j \in \{1, \dots, K\}, i \neq j} \left[ 1 - 6 \sum_m \frac{(r_{m(i)} - r_{m(j)})^2}{p(p^2 - 1)} \right]$ |
| Jaccard Distance (Jaccard, 1912) | $\frac{\vert \widehat{S}_{(i)} \cap \widehat{S}_{(j)}\vert }{\vert \widehat{S}_{(i)} \cup \widehat{S}_{(j)}\vert }$ |

<sup>a</sup> $\hat{\boldsymbol{\mu}}$ : fitting procedure on the training set <sup>b</sup> $S_0$ : index of active set = $\{j; \beta_j^0 \neq 0\}$ <sup>c</sup> $\widehat{S}$ : index of the set of non-zero estimated coefficients = $\{j; \hat{\beta}_j \neq 0\}$ <sup>d</sup> $|A|$ : is the cardinality of set $A$

### Results

All reported results are based on 200 simulation runs. We graphically summarized the results across simulations 1-3 for model fit (Figure 5) and feature stability (Figure 6). The results for simulations 4-6 are shown in the **Supplemental Section B, Figures S1-S6**. We restrict our attention to *SNR* =1, *ρ* = 0.9, and *α*_*j*_ ~ Unif [1.9, 2.1]. The model names are labeled as `summary measure_model` (e.g. `avg_lasso` corresponds using the average of the features in a cluster as inputs into a lasso regression model). When there is no summary measure appearing in the model name, that indicates that the original variables were used (e.g. `enet` means all separate features were used in the elasticnet model). In panel A of Figure 5, we plot the true positive rate against the false positive rate for each of the 200 simulations. We see that across all simulation scenarios, the SEPARATE method has extremely poor sensitivity compared to both CLUST and ECLUST, which do much better at identifying the active variables, though the resulting models are not always sparse. The relatively few number of green points in panel A is due to the small number of estimated clusters (**Supplemental Section C, Figure S7**) leading to very little variability in performance across simulations. The better performance of ECLUST over CLUST is noticeable as more points lie in the top left part of the plot. The horizontal banding in panel A reflects the stability of the TOM-based clustering approach. ECLUST also does better than CLUST in correctly determining whether a feature is zero or nonzero (Figure 5, panel B). Importantly, across all three simulation scenarios, ECLUST outperforms the competing methods in terms of RMSE (Figure 5, panel C), regardless of the summary measure and modeling procedure. **We present the distribution for the effective number of variables selected in the supplemental material (Figures S8 and S9). We see that their distributions are similar, and in most cases, the median number of variables selected from ECLUST is less than the median number of variables selected from CLUST.**

**Figure 5.**
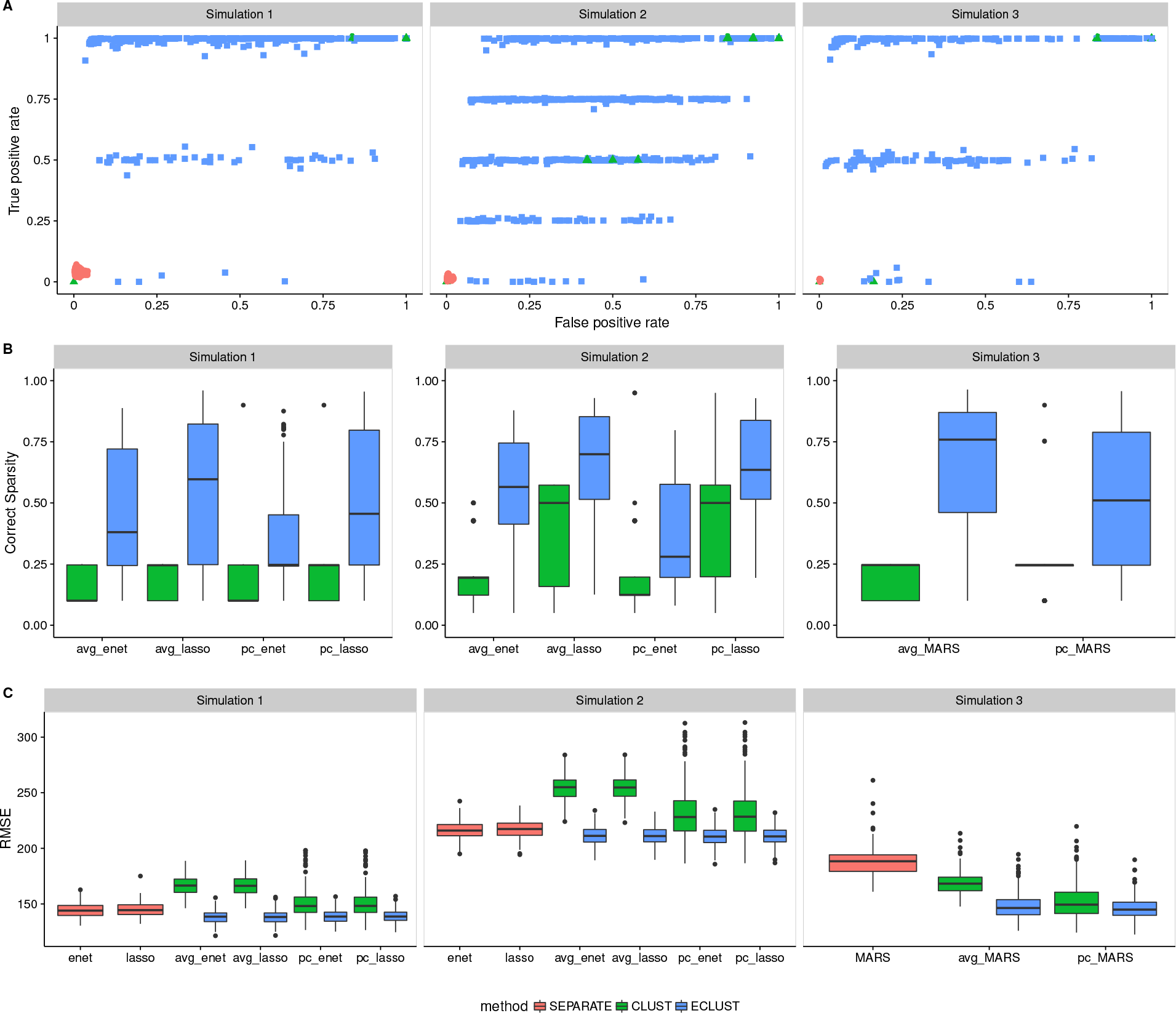
Model fit results from simulations 1, 2 and 3 with *SNR* =1, *ρ* = 0.9, and *α*_*j*_ ~ Unif [1.9, 2.1]. SEPARATE results are in pink, CLUST in green and ECLUST in blue.

**Figure 6.**
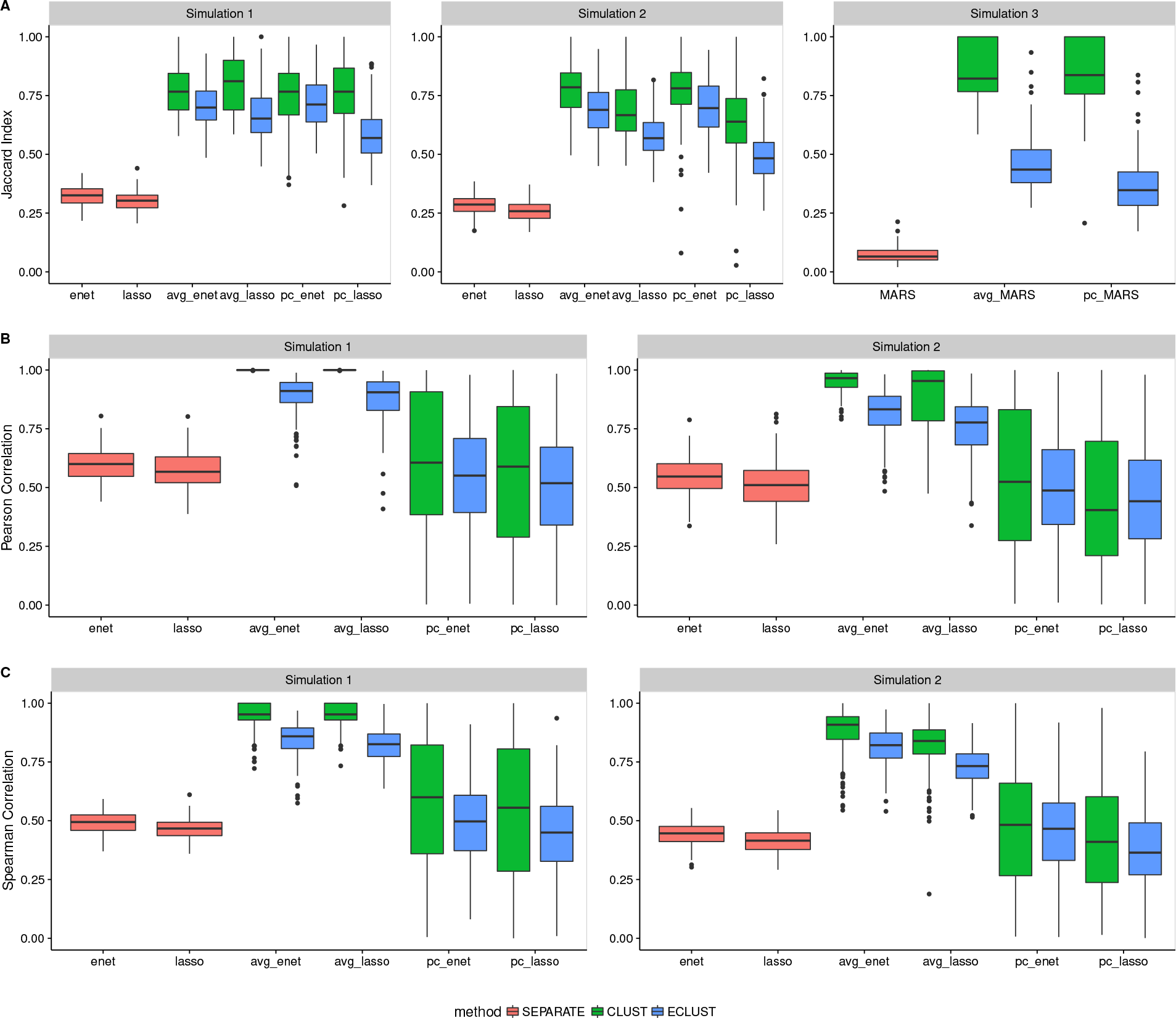
Stability results from simulations 1, 2 and 3 for *SNR* =1, *ρ* = 0.9, and *α*_*j*_ ~ Unif [1.9, 2.1]. SEPARATE results are in pink, CLUST in green and ECLUST in blue.

While the approach using all separate original variables (SEPARATE) produce sparse models, they are sensitive to small perturbations of the data across all stability measures (Figure 6), i.e, similar datasets produce very different models. Although the median for the CLUST approach is always slightly better than the median for ECLUST across all stability measures, CLUST results can be much more variable, particularly when stability is measured by the agreement between the value and the ranking of the estimated coefficients across CV folds (Figure 6, panels B and C). The number of estimated clusters, and therefore the number of features in the regression model, tends to be much smaller in CLUST compared to ECLUST, and this explains its poorer performance using the stability measures in Figure 6, since there are more coefficients to estimate. Overall, we observe that the relative performance of ECLUST versus CLUST in terms of stability is consistent across the two summary measures (average or principal component) and across the penalization procedures. **The complete results for different values of** *ρ*, *SNR* **and** *α*_*j*_ **(when applicable) are available in the Supplemental Section D, Figures S10-S15 for Simulation 1, Figures S16 - S21 for Simulation 2, and Figures S22 - S25 for Simulation 3. They** show that these conclusions are not sensitive to the *SNR*, *ρ* or *α*_*j*_. Similar conclusions are made for a binary outcome using logistic regression versions of the lasso, elasticnet and MARS. ECLUST and CLUST also have better calibration than the SEPARATE method for both linear and non-linear models (**Supplemental Section B, Figures S3-S6**). **The distributions of Hosmer-Lemeshow (HL)** *p***-values do not follow uniformity. This is in part due to the fact that the HL test has low power in the presence of continuous-dichotomous variable interactions (Hosmer et al., 1997). Upon inspection of the Q-Q plots, we see that the models have difficulty predicting risks at the boundaries which is a known issue in most models. We also have a small sample size of 200, which means there are on average only 20 subjects in each of the 10 bins. Furthermore, the HL test is sensitive to the choice of bins and method of computing quantiles. Nevertheless, the improved fit relative to the SEPARATE analysis is quite clear.**

We also ran all our simulations using the Pearson correlation matrix as a measure of similarity in order to compare its performance against the TOM. The complete results are in the **Supplemental Section E, Figures S26-S31 for Simulation 1, Figures S32 - S37 for Simulation 2, and Figures S38 - S41 for Simulation 3**. In general, we see slightly better performance of CLUST over ECLUST when using Pearson correlations. This result is probably due to the imprecision in the estimated correlations. The exposure dependent similarity matrices are quite noisy, and the variability is even larger when we examine the differences between two correlation matrices. Such large levels of variability have a negative impact on the clustering algorithm’s ability to detecting the true clusters.

## Analysis of three data sets

In this section we demonstrate the performance of ECLUST on three high dimensional datasets with contrasting motivations and features. In the first data set, normal brain development is examined in conjunction with intelligence scores. In the second data set we aim to identify molecular subtypes of ovarian cancer using gene expression data. The investigators’ goal in the third data set is to examine the impact of gestational diabetes mellitus (GDM) on childhood obesity in a sample of mother-child pairs from a prospective birth cohort. The datasets comprise a range of sample sizes, and both the amount of clustering in the HD data and the strength of the effects of the designated exposure variables vary substantially. Due to the complex nature of these datasets, we decided to use MARS models for step 2 of our algorithm for all 3 datasets, as outlined in Table 1. In order to assess performance in these data sets, we have computed the 0.632 estimator (Efron, 1983) and the 95% confidence interval of the *R*^2^ and RMSE from 100 bootstrap samples. The *R*^2^ reported here is defined as the squared Pearson correlation coefficient between the observed and predicted response (Kvålseth, 1985), and the RMSE is defined as in Table 3. Because MARS models can result in unstable predictors (Kuhn, 2008), we also report the results of bagged MARS from *B* = 50 bootstrap samples, where bagging (Breiman, 1996) refers to averaging the predictions from each of the MARS models fit on the *B* bootstrap samples.

### NIH MRI Study of Normal Brain Development

The NIH MRI Study of Normal Brain Development, started in 2001, was a 7 year longitudinal multi-site project that used magnetic resonance technologies to characterize brain maturation in 433 medically healthy, psychiatrically normal children aged 4.5-18 years (Evans et al., 2006). The goal of this study was to provide researchers with a representative and reliable source of healthy control subject data as a basis for understanding atypical brain development associated with a variety of developmental, neurological, and neuropsychiatric disorders affecting children and adults. Brain imaging data (e.g. cortical surface thickness, intra-cranial volume), behavioural measures (e.g. IQ scores, psychiatric interviews, behavioral ratings) and demographics (e.g. socioeconomic status) were collected at two year intervals for three time points and are publicly available upon request. Previous research using these data found that level of intelligence and age correlate with cortical thickness (Shaw et al., 2006; Khundrakpam et al., 2013), but to our knowledge no such relation between income and cortical thickness has been observed. We therefore used this data to see the performance of ECLUST in the presence (age) and absence (income) of an effect on the correlations in the HD data. We analyzed the 10,000 most variable regions on the cortical surface from brain scans corresponding to the first sampled time point only. We used binary age (166 age ≤ 11.3 and 172 > 11.3) and binary income (142 high and 133 low income) indicator as the environment variables and standardized IQ scores as the response. We identified 22 clusters from *TOM*(*X*_all_) and 57 clusters from *TOM*(*X*_diff_) when using age as the environment, and 86 clusters from *TOM*(*X*_all_) and 49 clusters from *TOM*(*X*_diff_) when using income as the environment. Results are shown in Figure 7, panels C and D. The method which uses all individual variables as predictors (pink), has better *R*^2^ but also worse RMSE compared to CLUST and ECLUST, likely due to over-fitting. There is a slight benefit in performance for ECLUST over CLUST when using age as the environment (panel D). Importantly, we observe very similar performance between CLUST and ECLUST across all models (panel C), suggesting very little impact on the prediction performance when including features derived both with and without the *E* variable, in a situation where they are unlikely to be relevant.

**Figure 7.**
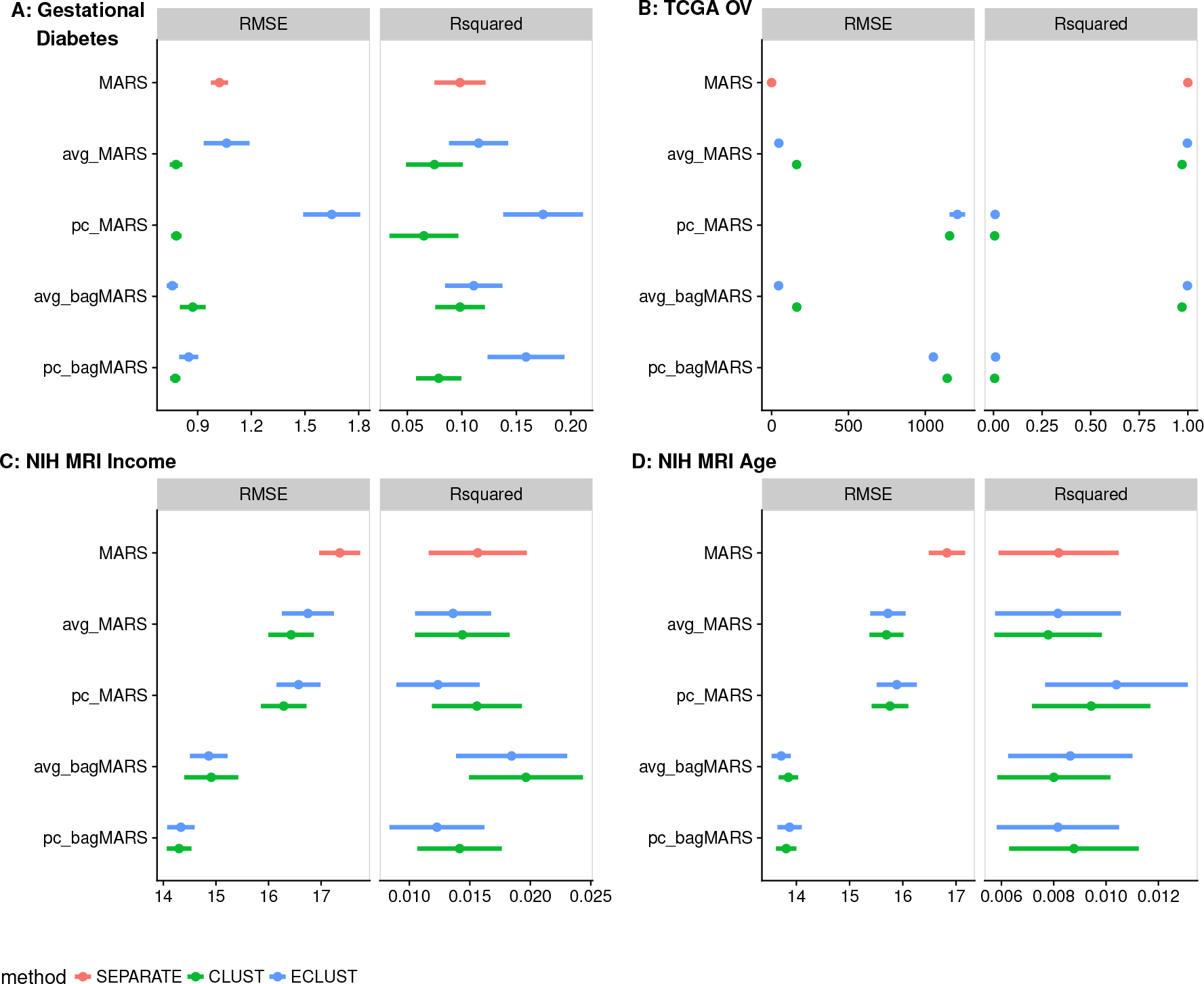
Model fit measures from analysis of three data sets: (A) Gestational diabetes birth-cohort (B) TCGA Ovarian Cancer study (C) NIH MRI Study with income as the environment variable (D) NIH MRI Study with age as the environment variable

### Gene Expression Study of Ovarian Cancer

Differences in gene expression profiles have led to the identification of robust molecular subtypes of ovarian cancer; these are of biological and clinical importance because they have been shown to correlate with overall survival (Tothill et al., 2008). Improving prediction of survival time based on gene expression signatures can lead to targeted therapeutic interventions (Helland et al., 2011). The proposed ECLUST algorithm was applied to gene expression data from 511 ovarian cancer patients profiled by the Affymetrix Human Genome U133A 2.0 Array. The data were obtained from the TCGA Research Network: http://cancergenome.nih.gov/ and downloaded via the TCGA2STAT R library (Wan et al., 2015). Using the 881 signature genes from Helland et al. (2011) we grouped subjects into two groups based on the results in this paper, to create a “positive control” environmental variable expected to have a strong effect. Specifically, we defined an environment variable in our framework as: *E* = 0 for subtypes C1 and C2 (*n* = 253), and *E* =1 for subtypes C4 and C5 (*n* = 258). Overall survival time (log transformed) was used as the response variable. Since these genes were ascertained on survival time, we expected the method using all genes without clustering to have the best performance, and hence one goal of this analysis was to see if ECLUST performed significantly worse as a result of summarizing the data into a lower dimension. We found 3 clusters from *TOM*(*X*_all_) and 3 clusters from *TOM*(*X*_diff_); results are shown in Figure 7, panel C. Across all models, ECLUST performs slightly better than CLUST. Furthermore it performs almost as well as the separate variable method, with the added advantage of dealing with a much smaller number of predictors (881 with SEPARATE compared to 6 with ECLUST).

### Gestational diabetes, epigenetics and metabolic disease

Events during pregnancy are suspected to play a role in childhood obesity development but only little is known about the mechanisms involved. Indeed, children born to women who had GDM in pregnancy are more likely to be overweight and obese (Wendland et al., 2012), and evidence suggests epigenetic factors are important piece of the puzzle (Bouchard et al., 2010, 2012). Recently, methylation changes in placenta and cord blood were associated with GDM (Ruchat et al., 2013), and here we explore how these changes are associated with obesity in the children at the age of about 5 years old. DNA methylation in placenta was measured with the Infinium HumanMethylation450 BeadChip (Illumina, Inc (Bibikova et al., 2011)) microarray, in a sample of 28 women, 20 of whom had a GDM-affected pregnancy, and here, we used GDM status as our *E* variable, assuming that this has widespread effects on DNA methylation and on its correlation patterns. Our response, *Y*, is the standardized body mass index (BMI) in the offspring at the age of 5. In contrast to the previous two examples, here we had no particular expectation of how ECLUST would perform. Using the 10,000 most variable probes, we found 2 clusters from *TOM*(*X*_all_), and 75 clusters from *TOM*(*X*_diff_). The predictive model results from a MARS analysis are shown in Figure 7, panel A. When using *R*^2^ as the measure of performance, ECLUST outperforms both SEPARATE and CLUST methods. When using RMSE as the measure of model performance, performance tended to be better with CLUST rather than ECLUST perhaps in part due to the small number of clusters derived from *TOM*(*X*_all_) relative to *TOM*(*X*_diff_). Overall, the ECLUST algorithm with bagged MARS and the 1st PC of each cluster performed best, i.e., it had a better *R*^2^ than CLUST with comparable RMSE. The sample size here is very small, and therefore the stability of the model fits is limited stability. The probes in these clusters mapped to 164 genes and these genes were selected to conduct pathway analyses using the Ingenuity Pathway Analysis (IPA) software (Ingenuity System). IPA compares the selected genes to a reference list of genes included in many biological pathways using a hypergeometric test. Smaller *p* values are evidence for over-represented gene ontology categories in the input gene list. The results are summarized in Table 4 and provide some biological validation of our ECLUST method. For example, the Hepatic system is involved with the metabolism of glucose and lipids (Saltiel and Kahn, 2001), and behavior and neurodevelopment are associated with obesity (Epstein et al., 2004). Furthermore, it is interesting that embryonic and organ development pathways are involved since GDM is associated with macrosomia (Ehrenberg et al., 2004).

**Table 4.** Ingenuity Pathway Analysis Results - top-ranked diseases and disorders, and physiological system development and function epigentically affected by gestational diabetes mellitus and associated with childhood body mass index

| Category | Name | *p* values | *n*<sup>a</sup> |
| --- | --- | --- | --- |
| Diseases and Disorders | Hepatic System Disease | [9.61e−7 – 5.17e−7] | 75 |
| Physiological System Development and Function | Behavior | [1.35e−2 – 7.82e−8] | 33 |
|  | Embryonic Development | [1.35e−2 – 2.63e−8] | 26 |
|  | Nervous System Development and Function | [ 1.35e−2 – 2.63e−8] | 43 |
|  | Organ Development | [1.35e−2 – 2.63e−8] | 20 |
|  | Organismal Development | [1.35e−2 – 2.63e−8] | 34 |

<sup>a</sup>number of genes involved in each pathway

## Discussion

The challenge of precision medicine is to appropriately fit treatments or recommendations to each individual. Data such as gene expression, DNA methylation levels, or magnetic resonance imaging (MRI) signals are examples of HD measurements that capture multiple aspects of how a tissue is functioning. These data often show patterns associated with disease, and major investments are being made in the genomics research community to generate such HD data. Analytic tools increasing prediction accuracy are needed to maximize the productivity of these investments. However, the effects of exposures have usually been overlooked, but these are crucial since they can lead to ways to intervene. Hence, it is essential to have a clear understanding of how exposures modify HD measures, and how the combination leads to disease. Existing methods for prediction (of disease), that are based on HD data and interactions with exposures, fall far short of being able to obtain this clear understanding. Most methods have low power and poor interpretability, and furthermore, modelling and interpretation problems are exacerbated when there is interest in interactions. In general, power to estimate interactions is low, and the number of possible interactions could be enormous. Therefore, here we have proposed a strategy to leverage situations where a covariate (e.g. an exposure) has a wide-spread effect on one or more HD measures, e.g. GDM on methylation levels. We have shown that this expected pattern can be used to construct dimension-reduced predictor variables that inherently capture the systemic covariate effects. These dimension-reduced variables, constructed without using the phenotype, can then be used in predictive models of any type. In contrast to some common analysis strategies that model the effects of individual predictors on outcome, our approach makes a step towards a systems-based perspective that we believe will be more informative when exploring the factors that are associated with disease or a phenotype of interest. We have shown, through simulations and real data analysis, that incorporation of environmental factors into predictive models in a way that retains a high dimensional perspective can improve results and interpretation for both linear and non linear effects.

We proposed two key methodological steps necessary to maximize predictive model interpretability when using HD data and a binary exposure: (1) dimension reduction of HD data built on exposure sensitivity, and (2) implementation of penalized prediction models. In the first step, we proposed to identify exposure-sensitive HD pairs by contrasting the TOM between exposed and unexposed individuals; then we cluster the elements in these HD pairs to find exposure-sensitive co-regulated sets. New dimension-reduced variables that capture exposure-sensitive features (e.g. the first principal component of each cluster) were then defined. In the second step we implemented linear and non-linear variable selection methods using the dimension-reduced variables to ensure stability of the predictive model. The ECLUST method has been implemented in the `eclust` (Bhatnagar, 2017) R package publicly available on CRAN. Our method along with computationally efficient algorithms, allows for the analysis of up to 10,000 variables at a time on a laptop computer.

The methods that we have proposed here are currently only applicable when three data elements are available. Specifically a binary environmental exposure, a high dimensional dataset that can be affected by the exposure, and a single phenotype. When comparing the TOM and Pearson correlations as a measure of similarity, our simulations showed that the performance of ECLUST was worse with correlations. This speaks to the potential of developing a better measure than the difference of two matrices. For example, we are currently exploring ways in which to handle continuous exposures or multiple exposures. The best way to construct an exposure-sensitive distance matrix that can be used for clustering is not obvious in these situations. One possible solution relies on a non-parametric smoothing based approach where weighted correlations are calculated. These weights can be derived from a kernel-based summary of the exposure covariates (e.g. (Qiu et al., 2016)). Then, contrasting unweighted and weighted matrices will allow construction of covariate-sensitive clusters. The choice of summary measure for each cluster also warrants further study. While principal components and averages are well understood and easy to implement, the main shortcoming is that they involve all original variables in the group. As the size of the groups increase, the interpretability of these measures decreases. Non-negative matrix factorization (Lee and Seung, 2001) and sparse principal component analysis (SPCA) (Witten et al., 2009) are alternatives which find sparse and potentially interpretable factors. Furthermore, structured SPCA (Jenatton et al., 2009) goes beyond restricting the cardinality of the contributing factors by imposing some a priori structural constraints deemed relevant to model the data at hand.

We are all aware that our exposures and environments impact our health and risks of disease, however detecting how the environment acts is extremely difficult. Furthermore, it is very challenging to develop reliable and understandable ways of predicting the risk of disease in individuals, based on high dimensional data such as genomic or imaging measures, and this challenge is exacerbated when there are environmental exposures that lead to many subtle alterations in the genomic measurements. Hence, we have developed an algorithm and an easy-to use software package to transform analysis of how environmental exposures impact human health, through an innovative signal-extracting approach for high dimensional measurements. Evidently, the model fitting here is performed using existing methods; our goal is to illustrate the potential of improved dimension reduction in two-stage methods, in order to generate discussion and new perspectives. If such an approach can lead to more interpretable results that identify gene-environment interactions and their effects on diseases and traits, the resulting understanding of how exposures influence the high-volume measurements now available in precision medicine will have important implications for health management and drug discovery.

## Availability of data and material

1. NIH MRI Study of Normal Brain Development data are available in the Pediatric MRI Data Repository, https://pediatricmri.nih.gov/
2. Gene Expression Study of Ovarian Cancer data are available in the Genomic Data Commons repository, https://gdc.cancer.gov/, and were downloaded via the TCGA2STAT R library Wan et al. (2015)
3. Gestational diabetes, epigenetics and metabolic disease: the clinical data, similarity matrices and cluster summaries are available at Zenodo [10.5281/zenodo.259222]. The raw analysed during the current study are not publicly available due to reasons of confidentiality, although specific collaborations with LB can be requested.

## Competing interests

The authors declare that they have no competing interests.

## Author’s contributions

SRB, CMTG, YY, MB: conceptualization; SRB, LB, BK: data curation; SRB: formal analysis, software, visualization; SRB, CMTG: methodology; SRB, CMTG: writing-original draft; SRB, CMTG, YY, BK, ACE, MB, LB: writing-review & editing.

## Acknowledgements

SRB was supported by the Ludmer Centre for Neuroinformatics and Mental Health.

## Additional Files

**Additional file 1 — Supplemental Methods and Simulation Results**

Contains the following sections:

A. **Description of Topological Overlap Matrix** – detailed description of the TOM
B. **Binary Outcome Simulation Results** – results of simulations 4, 5 and 6
C. **Analysis of Clusters** – number of estimated clusters by different measures of similarity
D. **Simulation Results Using TOM as a Measure of Similarity** – detailed simulation results using TOM as a measure of similarity
E. **Simulation Results Using Pearson Correlations as a Measure of Similarity** – detailed simulation results using Pearson Correlations as a measure of similarity
F. **Visual Representation of Similarity Matrices** – similarity matrices based on Pearson’s correlation coefficient

